# Fidelity-Ensuring Consistency of Mitosis is Safeguarded by the 53BP1-USP28-p53 Pathway

**DOI:** 10.64898/2026.03.16.712228

**Authors:** Avital Shulman, Kanako Ozaki, Lukas R. Chang, Eden Hoong, Meng-Fu Bryan Tsou

## Abstract

Mitosis is monitored by and mechanically coupled to the spindle assembly checkpoint (SAC), which halts mitotic progression until fidelity is met, disregarding time efficiency. Conversely, there exists SAC-independent, efficiency-promoting surveillance mechanically uncoupled from mitosis. This external mitotic surveillance (EMS), comprising 53BP1, USP28, and p53, induces post-mitotic arrest following prolonged, inefficient mitosis. To explore additional EMS inputs, we performed comparative CRISPR-Cas9 screens for genes functionally safeguarded by EMS, identifying, among others, two components of the RZZ kinetochore complex—KNTC1 and ZWILCH. Depleting KNTC1, which impairs fidelity-ensuring activities of mitosis, including SAC, triggered population-level post-mitotic arrests without widespread, characteristic mitotic delay or catastrophe. Instead, KNTC1 depletion produced mostly viable mitosis and yet activated EMS via ectopic accumulation of 53BP1-USP28-p53 complexes over normal mitotic duration. These results suggest that when the fidelity-ensuring control within mitosis is itself compromised, mitosis is rendered invalid from without by EMS, echoing Gödel’s incompleteness theorems.

## Introduction

Mitosis is a tightly orchestrated process that ensures faithful chromosome segregation during cell division. To preserve genomic stability, cells must coordinate complex events such as G2-to-M entry, spindle formation, chromosome alignment, and timely anaphase onset [1–3]. The spindle assembly checkpoint (SAC) serves as a critical safeguard that halts anaphase onset until all chromosomes are properly attached to spindle microtubules [3, 4], mechanically coupling mitotic progression and duration to the fidelity of chromosome-spindle attachments. Mitosis as a fidelity-ensuring system, however, carries a logical vulnerability: what safeguards exist if the SAC is itself compromised? Like Gödel’s second incompleteness theorem positing that a system cannot prove its own consistency from within [5], cells may require separate surveillance from outside mitosis. Nevertheless, wild-type mitosis with optimal SAC typically proceeds efficiently within a narrow temporal window of ∼30 minutes, and deviations from this norm—particularly prolonged mitosis—can often be indicative of cellular distress. For example, SAC activation by persistent mitotic insults can hold mitosis for much extended durations (>10 hours), leading to mitotic slippage, DNA damage responses, and apoptosis [6–8].

Interestingly, recent studies have uncovered a separate surveillance mechanism, which is not part of SAC-mediated chromosome-spindle attachment fidelity nor mechanically coupled to mitotic progression, but it responds to mild extensions of mitotic duration and induces cell cycle arrest, even if mitosis completes successfully without DNA damage [9–11]. This external mitotic surveillance, here called EMS, composed of 53BP1, USP28, and p53, triggers a p21-mediated cell cycle arrest following mitoses that exceed a defined temporal threshold, regardless of whether SAC is present or not [9]. With or without EMS, mitosis per se proceeds unchanged under otherwise the same conditions. In the presence of EMS, human non-transformed RPE1 cells respond to ∼3-fold extensions in mitosis (>90 min) [10, 11] or repeated milder delays (60-90 min) [12] by assembling 53BP1–USP28–p53 complexes [12, 13]. These protein complexes accumulate during mitosis in a time-dependent manner, transmit into the next cell cycle as a form of memory, and induce cell proliferation arrest [12]. 53BP1 is known to mediate DNA damage response (DDR) during interphase [14] and localizes to nascent kinetochores in early mitosis [15], but neither the DDR activity [9] nor its kinetochore association [13] is required for EMS-mediated cell cycle arrest.

Through EMS, disruption of any cellular components or genes that either directly or indirectly causes mitotic delay can in theory generate the same terminal phenotype, post-mitotic arrest, adding complications to functional characterization and interpretation. Indeed, EMS was first elucidated through studies not on mitosis per se but on the ciliogenesis organelle, the centrosome, which is not essential for mitosis [16–19]. This raises a question of whether there are other genes or cellular elements whose functions are masked or safeguarded by EMS. To explore these questions, we conducted genome-wide, comparative CRISPR-Cas9 screens in wild-type (WT), *p53^-/-^*, and *USP28^-/-^* RPE1 cells to identify genes functionally masked by EMS. Among the top hits from our screens were KNTC1 (ROD) and ZWILCH, two core components of the ROD–ZW10–ZWILCH (RZZ) complex [20–22]. RZZ complexes assemble onto nascent kinetochores during early mitosis to promote outer kinetochore expansion and maturation [23, 24], enabling optimal SAC signaling and microtubule binding for fidelity [25–31]. Our studies revealed that loss of KNTC1, which impairs SAC maintenance—and thus overall integrity of mitosis—activates EMS by inducing ectopic formation of 53BP1–USP28–p53 complexes without characteristic mitotic delays, suggesting that fidelity of mitosis is protected by nested layers of surveillance, SAC and EMS.

## Results

### Loss of KNTC1 triggers a p53, USP28, and 53BP1 dependent cell-cycle arrest

#### CRISPR screens identify the RZZ complex as a novel cellular component selectively required for WT but not p53^-/-^ and USP28^-/-^ cells

Full genome, comparative CRISPR screens (Fig. 1A) were conducted to look for genes whose losses negatively impact WT cells but are near neutral or positive in *p53^-/-^* and *USP28^-/-^*backgrounds. Raw data of the screen are publicly available on GEO, ID # GSE306237 (<u>GEO link</u>). The neutrality cutoff value in our data analysis is defined and normalized in each screen using the "gold-standard" set of 927 non-essential genes [32] (see Methods). A list of 79 genes was identified as having two or more gRNAs significantly impacting the growth of WT but not *p53^-/-^* and *USP28^-/-^* cells (Table S1). Gene Ontology (GO) Enrichment Analysis for Cellular Components [33, 34] revealed two out of three components of the RZZ complex—KNTC1 (also known as ROD) and ZWILCH—as high-confidence hits, with >100-fold enrichment (Fig. 1B). Interestingly, centrosome assembly factors were not in the list (Table S1), as their losses, while producing prolonged mitosis viable in the absence of EMS, hindered the proliferation of *p53^-/-^* and *USP28^-/-^* cells significantly enough for them to be excluded, implying functional differences from our hits.

**Fig. 1:**
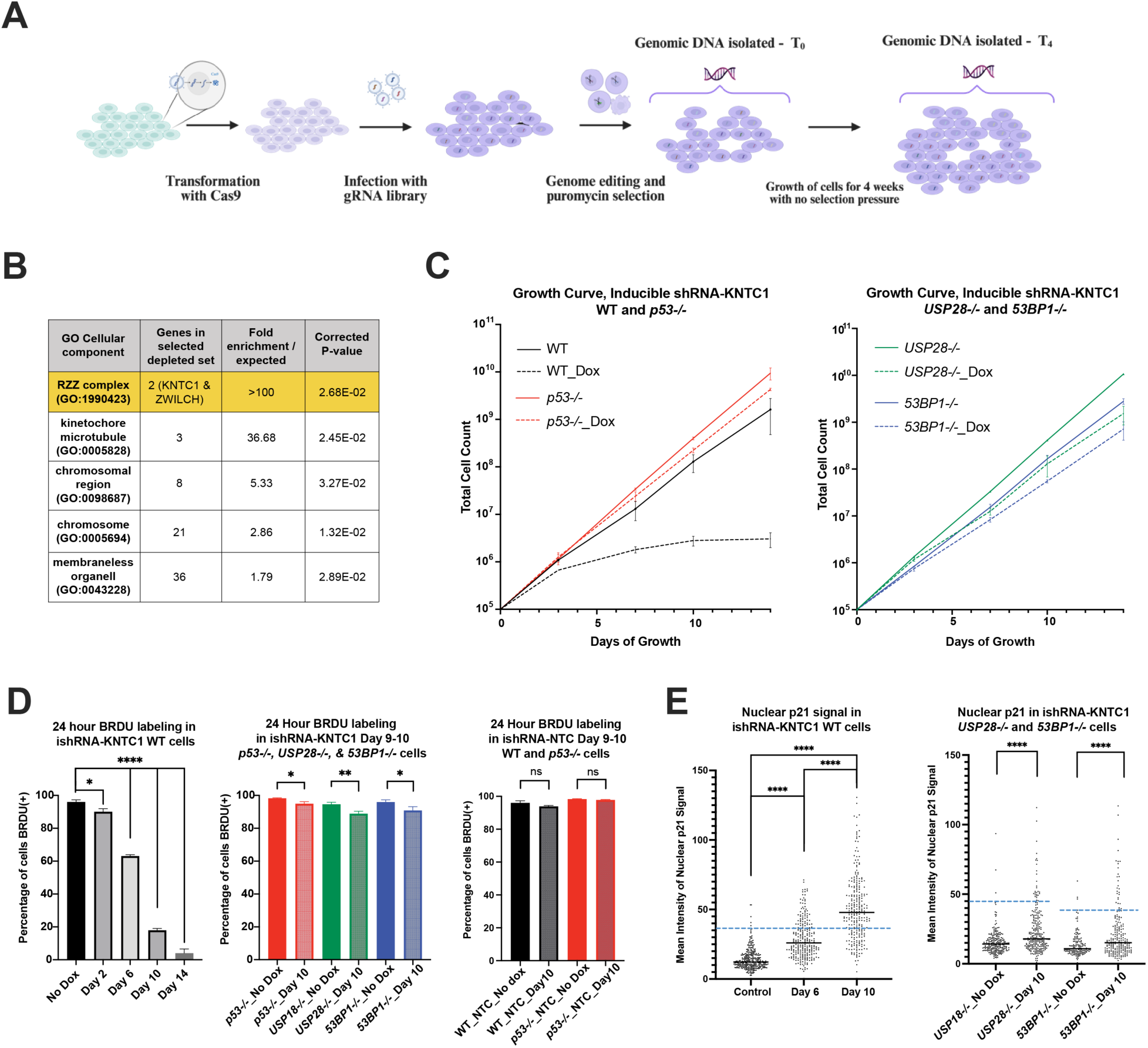
Loss of KNTC1 triggers a p53, USP28, and 53BP1 dependent cell-cycle arrest. **(A)** Schematic of full genome CRISPR screens. After four weeks of growth with no selection pressure, gRNAs selectively lost in WT but not *p53^-/-^* and *USP28^-/-^* cells were used to identify genes whose functions are guarded or masked by EMS. **(B)** GO Cellular Component Enrichment Analysis on the 79 genes that were considered essential in the WT but neutral or near neutral in *p53^-/-^*and *USP28^-/-^* backgrounds. The RZZ complex was revealed as a top hit, with two of three components, KNTC1 (ROD) and ZWILCH, identified. **(C)** Growth curve analyses showed that following induction of shRNA-KNTC1, WT (black) cells were fully arrested, while *p53^-/-^*(red), *USP28^-/-^* (green), and *53BP1^-/-^* (blue) cells continued to grow, albeit at slightly slower rates compared to controls (the respective shRNA clonal line without Dox induction). Data are mean ± SD, *N* = 3, initial cell count = 100,000, and doxycycline (Dox) at 1 μg/mL. **(D)** BrdU labeling assays revealed that KNTC1 knockdown has a widespread, detrimental growth impact on WT cells (left) but not *p53^-/-^, USP28-/-,* and *53BP1^-/-^* cells (center), compared to the non-targeting shRNA control (shRNA-NTC, right). Note that KNTC1 knockdown does have a non-population-level growth impact on EMS-knockout cells (*p53^-/-^, USP28^-/-^,* or *53BP1^-/-^*), consistent with (C). Cells were fixed at the end of indicated days, with BrdU (10 μM) added 24 hours prior to fixation. Data are mean ± SD, *n* > 300, *N* = 3, statistical significance calculated via T-test with Welch’s correction. **(E)** Quantification of nuclear p21 intensity at indicated days following knockdown of KNTC1 in WT (left), *USP28^-/-^* (right), or *53BP1^-/-^* (right) cells. At Day 10 after KNTC1 knockdown, ∼70% of WT cells showed a nuclear p21 intensity that is 3 standard deviations above the mean in control cells, which are defined as high outliers (above dashed blue lines). In contrast, only 10-11% of *USP28^-/-^* or *53BP1^-/-^* cells were high outliers by Day 10, consistent with the observation that KNTC1 loss has a mild, non-population-level growth impact on EMS-knockout cells. Data are mean signal intensity, *n* > 240. Black lines indicating median, dashed blue line marking 3 standard deviations above the control mean, and statistical significance calculated via T-test with Welch’s correction.

To confirm our screens, KNTC1 was knocked down by inducible mirE shRNA [35, 36] using simultaneous expression of two constructs targeting different coding exons of KNTC1 (exons 33 and 55). Consistent with our screen, growth curve analyses following shRNA induction showed arrest of WT cells, while *p53^-/-^*, *USP28^-/-^*, and *53BP1^-/-^* cells continue to grow, albeit at a slightly lower rate (Fig. 1C). Furthermore, 24-hour BrdU labeling assays showed significant decreases in the percentage of WT cells entering the cell cycle over time after shRNA induction (Fig. 1D, first panel). In contrast, a majority of *p53^-/-^*, *USP28^-/-^*, or *53BP1^-/-^* cells incorporated BrdU at Day 9-10 following shRNA induction (Fig. 1D, middle panel). No significant difference in BrdU incorporation is seen at Day 9-10 following induction of non-targeting shRNA control constructs in either WT or *p53^-/-^* cells (Fig. 1D, third panel), indicating that the arrest seen in WT cells is a specific response to the loss of KNTC1, rather than a stress response to shRNA expression.

Prolonged mitosis is known to activate EMS without DNA damage responses [9], causing p21-mediated cell cycle arrest as the terminal phenotype. We examined if loss of KNTC1 causes similar impacts. Consistently, a significant increase in nuclear accumulation of p21 was seen in WT cells following KNTC1 knockdown, with ∼70% of cells at Day 10 identified as high outliers, having a nuclear p21 signal that is at least 3 standard deviations above the mean signal in control cells (Fig. 1E, first panel, outliers above the dashed blue line). In contrast, in *USP28^-/-^* or *53BP1^-/-^* cells at Day 10 following KNTC1 knockdown (Fig. 1E, second panel, outliers above respective dashed blue lines), only ∼11% of cells have a nuclear p21 signal considered a high outlier to the controls. This is consistent with our growth curve and BrdU incorporation assays, showing that following KNTC1 loss in *USP28^-/-^*and *53BP1^-/-^* backgrounds, cell proliferation continues at the population level, with a slight increased background rate of cellular arrest. Moreover, using γH2AX foci as the marker, no elevated DNA damage responses following KNTC1 knockdown were observed (Fig. S1A-C). Together these results suggest that loss of KNTC1, like prolonged mitosis, induces EMS activation but not DNA damage responses, resulting in a population-level p21 upregulation and cell cycle arrest downstream of 53BP1 and USP28.

#### Both homozygous and heterozygous inactivation of KNTC1 can induce selective vulnerability in WT but not p53^-/-^, USP28^-/-^, and 53BP1^-/-^ cells

We next tested if a stable KNTC1 knockout could be generated in *p53^-/-^*cells using CRISPR/Cas9 gene editing. 5 days after infection with lentiviruses carrying an sgRNA targeting KNTC1, infected cells were individually seeded onto 96-well plates for clonal cell propagation and isolation. 35 growing clones were randomly selected, examined for KNTC1 localization at kinetochores at the population level, and in turn classified either as KNTC1-normal or KNTC1-disrupted clones (Fig. 2A). Under the *p53^-/-^* background, which is expected to tolerate KNTC1 loss, 4 out of 35 clones examined showed bright kinetochore KNTC1 as normally seen in WT (KNTC1^+/+^) cells, whereas the remainder showed disrupted staining, with weaker or no identifiable KNTC1 puncta at the kinetochores (Fig. 2A & 2B). These patterns suggest that perhaps even inactivating one allele of the KNTC1 gene in RPE1 cells, here called heterozygous knockout, can result in significant KNTC1 reduction at kinetochores. Indeed, sequencing the genomic cut site confirmed 2 randomly selected clones with no kinetochore KNTC1 signal as homozygous knockout and the other 2 with reduced KNTC1 signals as heterozygous knockout (Figs. 2A & 2B).

**Fig. 2.**
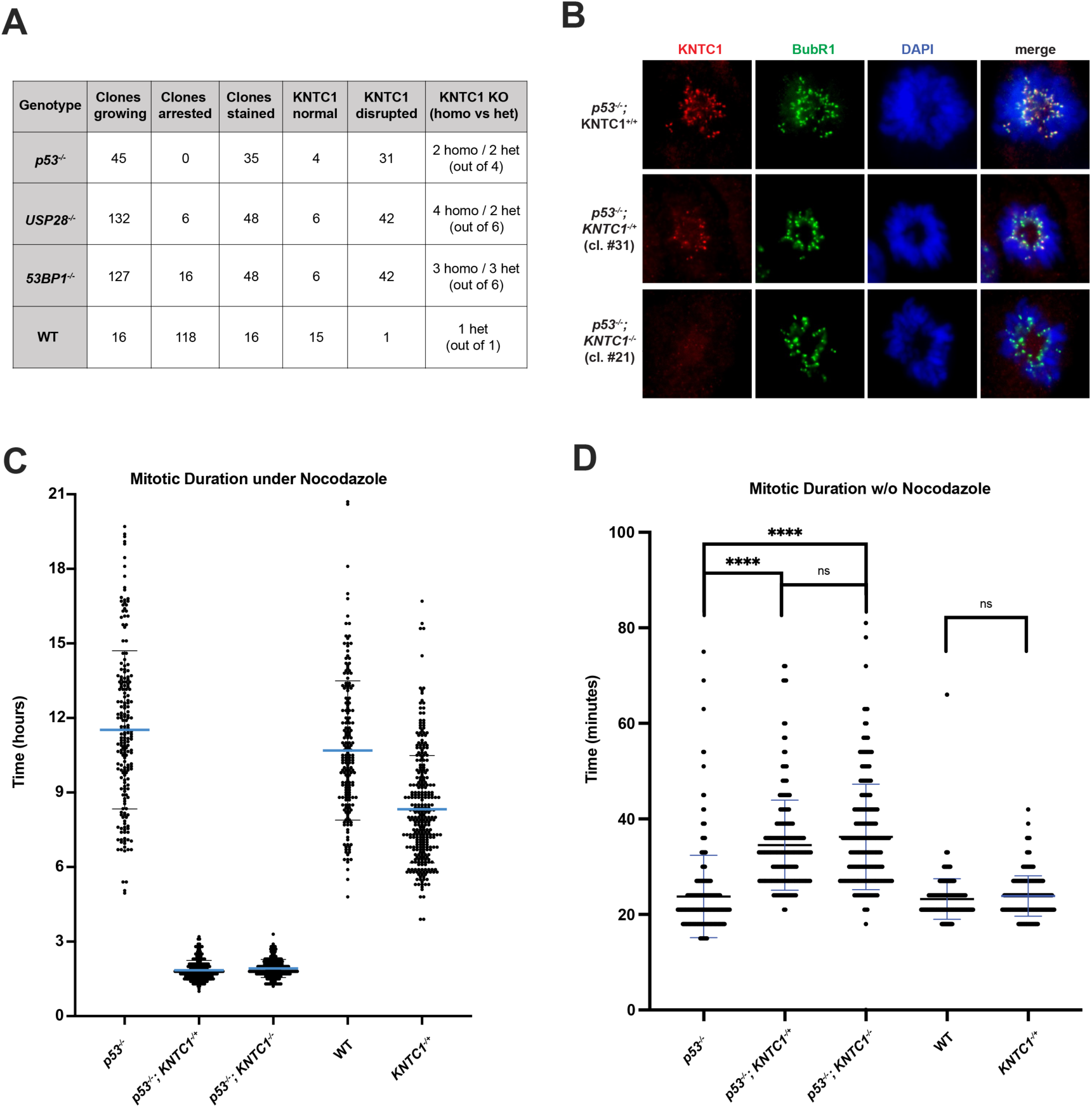
Both homozygous and heterozygous knockouts of KNTC1 can disrupt SAC and selectively arrest cells carrying intact EMS. **(A)** Analyses of clonal isolations for cells undergoing CRISPR-mediated KNTC1 targeting in *p53^-/-^*, *USP28^-/-^*, *53BP1^-/-^*, and WT backgrounds. The table starts with the total number of clones examined, among which how many were visibly arrested. Also tabulated are numbers of clones checked for KNTC1 staining at early mitotic kinetochores, of which how many exhibited normal or disrupted (reduced or absent) signals. Indicated numbers of clones with disrupted signals were further selected for sequencing, determining if they are heterozygous or homozygous KNTC1 knockouts. **(B)** Representative images of normal, reduced, and absent KNTC1 signals in *p53^-/-^; KNTC1^+/+^* (WT), *p53^-/-^; KNTC1^+/-^* (heterozygous knockout), and *p53^-/-^; KNTC1^-/-^* (homozygous knockout), respectively. **(C)** Quantification of mitotic durations prior to slippage in the presence of nocodazole. KNTC1^+/+^ cells in both *p53^-/-^* and WT backgrounds could arrest in nocodazole-treated mitosis for a mean duration of 11.5 and 10.7 hours, respectively, before slippage. In contrast, both *KNTC1^+/-^* (heterozygous) and *KNTC1^-/-^* (homozygous) cells in the *p53^-/-^* background exited mitosis after a short arrest of 1.8 hours, indicating a severe disruption of SAC. The singular *KNTC1^+/-^* clone (Singleton-Het) survived in the WT background, however, could stay in mitotic arrest for a mean of 8.3 hours, indicating a mildly compromised SAC. *n* > 200, nocodazole added at 50 ng/mL. **(D)** Quantification of normal mitotic duration in the absence of nocodazole perturbation. *KNTC1^+/-^* and *KNTC1^-/-^* cells obtained from the *p53^-/-^* background showed a small increase in average mitotic duration from 23 to 34 and 36 minutes, respectively, suggesting the presence of sub-optimal kinetochore function. The singular heterozygous KNTC1 clone survived in the WT background in our screen showed no significant increase in mitotic duration, remaining at 24 minutes, suggesting the presence of functional kinetochores or SAC. *n > 200,* statistical significance calculated via T-test with Welch’s correction.

To see if heterozygous knockout of KNTC1 can impact cell proliferation via EMS activation, clonal cell isolations following CRISPR/Cas9 gene editing were conducted in *USP28^-/-^*, *53BP1^-/-^*, and WT backgrounds (Fig. 2A). In each case, among the total number of clones identifiable at 3 weeks of growth across 96 well plates, the number of those that appeared visibly arrested with a small cluster of <10 senescent-like cells were counted. In the *USP28^-/-^* background, 138 total clones were identified at 3 weeks, with 6 visibly arrested, whereas in the *53BP1^-/-^* background, 16 out of 143 clones were visibly arrested (Fig. 2A). In both cases, KNTC1 localization at kinetochores was examined in 48 growing clones at random, finding 6 out of 48 to be KNTC1-normal and 42 to be KNTC1-disrupted. Furthermore, sequencing 6 KNTC1-disrupted clones from each population revealed the presence of both homozygous and heterozygous knockout clones (Fig. 2A; Fig. S2A, rows 1-6).

In striking contrast, clonal isolation following KNTC1 gene targeting in the WT background resulted in 134 clones identified, of which 118 were visibly arrested and only 16 were actively growing. After examining all 16 growing clones for kinetochore KNTC1, 15 were identified as KNTC1-normal and 1 as KNTC1-disrupted (Fig. 2A; Fig. S2A, bottom row), which we confirmed by sequencing to be a heterozygous knockout (Fig. 2A). The ratio of 15 out of 134 clones having normal KNTC1 signals is consistent with the rate of KNTC1-normal clones identified in other backgrounds, or the rate of expected WT clones following CRISPR-mediated gene targeting. However, the observation that only 1 KNTC1-disrupted clone, out of the expected ∼60, survived the clonal isolation suggests that heterozygous knockout of KNTC1 can widely induce cell cycle arrest in the presence of intact EMS.

#### Heterozygous knockout of KNTC1 can disrupt SAC severely enough to induce cell cycle arrest

As heterozygous knockout of KNTC1 induces EMS-dependent cell cycle arrest in most cases, we further examined if it has similar functional impacts on SAC integrity like homozygous knockouts. Time-lapse microscopy of cells growing in the presence of nocodazole—which disrupts kinetochore-spindle attachments—was conducted to measure how long cells can hold in mitosis before slipping into tetraploid G1. We examined the parental *p53^-/^*^-^ clone and its derived clones that are either heterozygous or homozygous for KNTC1 knockout, as well as the WT p53 clone and its surviving singleton carrying heterozygous knockout of KNTC1—p53^+/+^*; KNTC1^-/+^*cells—referred to hereafter as "Singleton-Het." Analysis showed that *p53^-/-^;* KNTC1^+/+^ cells stayed arrested in mitosis for a mean of 11.5 hours, with a range from 4.9 to 19.7 hours, while both *p53^-/-^; KNTC1^-/+^* and *p53^-/-^; KNTC1^-/-^* cells held in mitosis for significantly reduced durations, with means of 1.8 and 1.9 hours, ranging from 1.0 to 3.2 and 1.2 to 3.3 hours, respectively (Fig. 2C; Movies S1-3). In contrast, for the Singleton-Het that survives the clonal isolation, the mean mitotic duration under nocodazole dropped mildly from 10.7 hours seen in WT cells to 8.3 hours (Fig. 2C; Movies S4-5), suggesting that SAC activities are relatively less disrupted in this surviving p53^+/+^*; KNTC1^-/+^* singleton.

We next examined the average duration of mitosis in the above clones without nocodazole perturbation, finding that the mean duration increased from 24 minutes in *p53^-/-^; KNTC1^+/+^* cells to 34 or 36 minutes in the *p53^-/-^; KNTC1^-/+^* or *p53^-/-^; KNTC1^-/-^* cells, respectively (Fig. 2D), suggesting that heterozygous loss of KNTC1 can cause a functional disruption significant enough to impair SAC maintenance (Fig. 2C) while mildly increasing average mitotic duration. In contrast, for the surviving Singleton-Het, the average mitotic duration was unchanged, showing no significant difference from that of the control or wild-type cells (Fig. 2D), correlating with a lesser degree of functional disruption to SAC or kinetochores. Taken together, our results suggest that in most cases heterozygous loss of KNTC1 results in functional disruption significant enough to trigger EMS activation and post-mitotic arrest, resulting in loss of heterozygous clones in the WT background.

### Cell cycle arrest following the loss of KNTC1 does not require prolonged mitotic delays

#### Mitotic durations following the loss of KNTC1 are below the threshold needed for canonical EMS activation

To determine if cell cycle arrest is always preceded by longer mitoses known to activate EMS, mitotic durations of individual cells were tracked for multiple generations following acute, inducible KNTC1 depletion. We first determined the temporal threshold for EMS activation in our wild-type RPE1 ishRNA-KNTC1 cell line and found that individual mitotic divisions longer than 100-110 minutes effectively trigger cell cycle arrest (Fig. 3A). We then used time-lapse microscopy to examine the length of each division after induction of KNTC1 knockdown, with cells being filmed for 8 days starting on Day 2—48 hours after shRNA induction—through the end of Day 9. We found that following shRNA induction there is an increase in the range of mitotic durations, with scattered divisions observed up to 78 and down to 21 minutes in length (Fig. 3B, first panel). However, over 90% of divisions were completed in under 50 minutes, and the mean duration increased only slightly to 36 minutes, consistent with the average delays seen in stable knockout clones but substantially lower than the temporal threshold of 100-110 minutes for mitotic delay to induce arrest in this cell line.

**Fig. 3:**
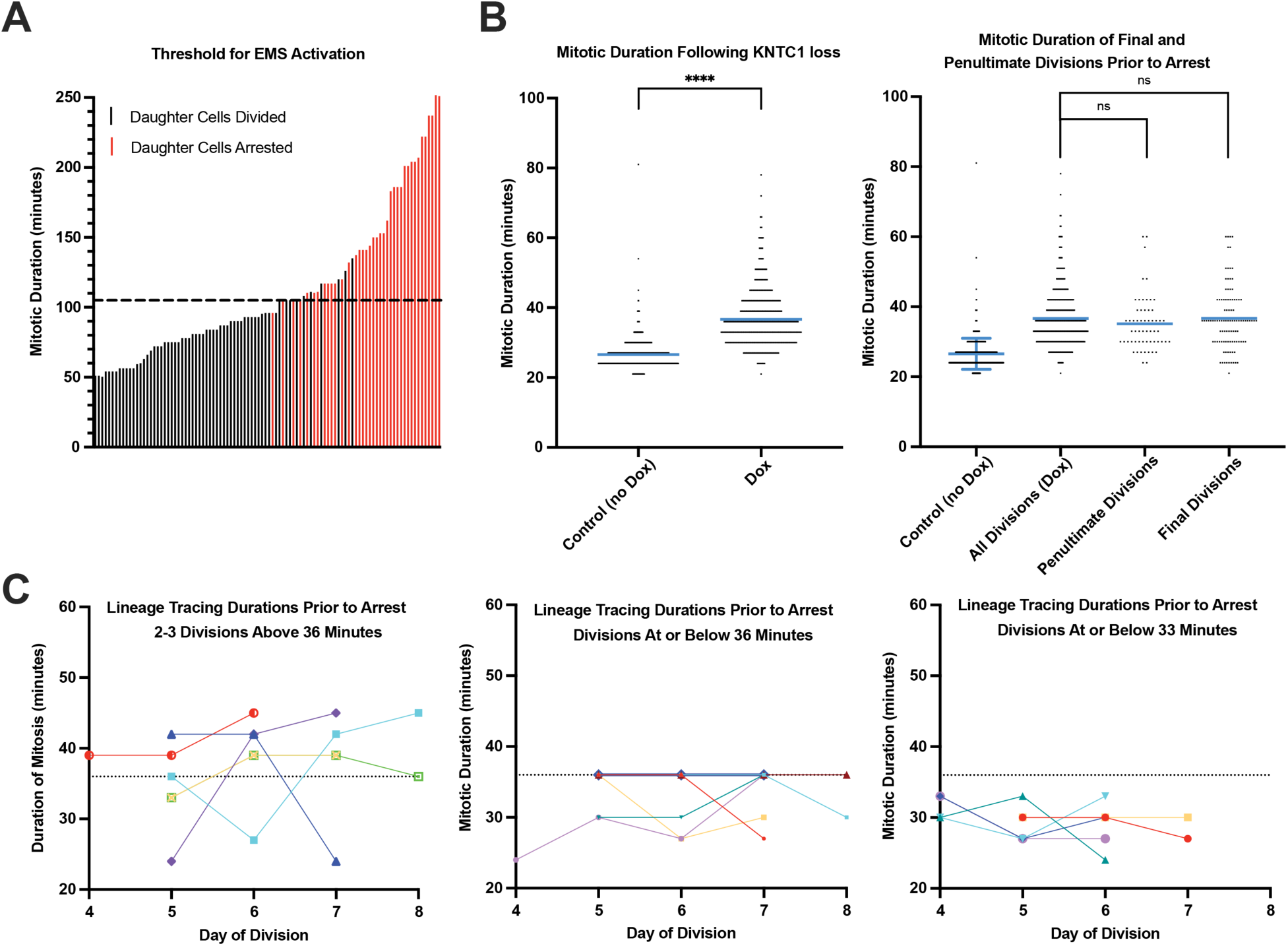
Loss of KNTC1 activates EMS without widespread, characteristic prolonged mitosis. **(A)** Identification of the mitotic duration threshold for EMS activation and cell cycle arrest in clonal ishRNA-KNTC1 cells. Time-lapse microscopy movies of asynchronously growing cells treated with nocodazole for 5 hours and then washed out of the drug for additional 50 hours were analyzed, revealing 105-110 minutes as the duration threshold. Dashed lines mark 105 minutes. **(B)** Examining the gradual impact of inducible KNTC1 knockdown on mitotic duration. Left, mitotic durations determined by filming ishRNA-KNTC1 cells for 10 days, during which KNTC1 knockdown was induced (Dox) or not induced (Control). Measurements of mitotic duration that were included in the plots started at the end of Day 2 (48 hrs after Dox) and ended at Day 10. Mitotic durations following the loss of KNTC1 were found to increase from a mean of 27 minutes to 36 minutes, with scattered divisions up to 80 minutes, all of which are lower than the EMS activation threshold of 100-110 minutes. *n* > 120, *N =3*, statistical analysis showed no difference between the datasets obtained in the separate experiments (not shown), and data are combined for total *n* > 450 divisions in both the control and Dox populations. Right, lineage tracking of sequential divisions revealed no difference in average durations of penultimate or final divisions prior to arrest, indicating that arrest is not due to repeated divisions above a threshold higher than the mean duration. *n* > 50 and *n* > 100 penultimate and final divisions, respectively, were identified within the Dox population. Blue lines at mean durations, ± SD indicated on the control population in the right panel. Statistical significances calculated via T-tests with Welch’s correction. **(C)** Of the 40 lineages with 3 or more divisions traced prior to arrest, only one was identified with three divisions above the mean of 36 minutes (left, red), and *n* = 5 with two divisions above the mean (left, assorted colors). *n* =13 and *n* = 6 instances were identified of 3-4 sequential divisions prior to arrest with durations at or below 36 and 33 minutes, respectively (center and right), indicating arrest can occur after repeated divisions with durations at/below the mean, and furthermore even within the mean + SD of the control population (highlighted in B, right panel).

Given the recent findings showing EMS activation via the cumulative effect of multiple mitotic divisions between 60-90 minutes [12], we asked if the arrest caused by KNTC1 depletion could be triggered by repeated milder delays in mitosis. To test this, we tracked cells across sequential divisions across the length of our time-lapse film following shRNA induction, noting the length of both the final and penultimate divisions prior to the arrest, which we defined as >60 hours with no divisions. Examining the distribution of these divisions (Fig. 3B, second panel) we found no significant difference in the overall distribution, nor in the mean duration—which remains at 36 minutes—of the final or penultimate divisions relative to all divisions observed under shRNA induction. Of the *n* =40 lineages in which 3 or more sequential divisions prior to arrest could be traced, only a single lineage was identified with three divisions above the mean of 36 minutes, and *n* = 5 were identified with two divisions above 36 minutes (Fig. 3C, first panel). More notably, *n* = 13 instances were identified of cells undergoing 3 or more divisions prior to arrest with durations at or below the mean of 36 minutes (Fig. 3C, center and right), of which *n* = 6 had 3 consecutive divisions below 33 minutes (Fig. 3C, right panel). Note that below 33 minutes is within the range of mitotic durations (the mean + standard deviation) seen in the control population (highlighted on the control population in the second panel of Fig. 3B). Thus, KNTC1-deficient cells can go through multiple, consecutive divisions with normal durations before arrest, with the last mitosis progressing no differently from the previous. These results together suggest that loss of KNTC1 can activate EMS without canonically associated prolonged mitotic delays.

#### EMS activation induced by KNTC1 loss does not require kinetochore localization of 53BP1

Kinetochore-associated 53BP1 is not required for EMS activation induced by prolonged mitosis [13]. However, the finding that EMS is activated without the characteristic prolonged mitotic delay, in response to the loss of a kinetochore complex, raised an idea that perhaps kinetochore localization of 53BP1 is involved in this case. Examining 53BP1 in *KNTC1^-/-^*cells revealed that it remains capable of localizing to kinetochores with no detectable difference (Fig. S3A, quantification not shown). We next sought to disrupt kinetochore 53BP1 for functional assays. It has been shown that 53BP1 localizes to kinetochores via its binding partner CENP-F and that a CENP-F point mutant (*CENP-F^E564P^*) selectively abolishes kinetochore recruitment of 53BP1 without affecting other known kinetochore functions [13]. RPE1 *CENP-F^E564P^* mutant cells were obtained [13], and examining these cells revealed that while 53BP1 is absent from kinetochores (Fig. S3B), kinetochore KNTC1 is unaffected (Fig. S3C). Moreover, *CENP-F^E564P^*cells could actively respond to prolonged mitotic delays as shown previously (data not shown). We then asked if knocking down KNTC1 via shRNA can induce cell cycle arrest in the *CENP-F^E564P^* background where kinetochore 53BP1 is lost. Growth curve analysis and BrdU incorporation following shRNA induction showed arrest of these cells, which could be rescued by full knockout of 53BP1 (Fig. S3D & E). Consistently, quantification of nuclear p21 intensity showed upregulation following shRNA induction in the *CENP-F^E564P^* background, rescued in *CENP-F^E564P^;53BP1^-/-^* (Fig. S3F). These results indicate that kinetochore localization of 53BP1 is not required for EMS activation in response to KNTC1 loss, nor to prolonged mitotic delay previously reported, leaving the functional purpose of kinetochore 53BP1 still unknown.

### Accumulation of 53BP1-USP28-p53 complexes in the absence of KNTC1 occurs soon after mitotic entry

We next performed the proximity ligation assay (PLA) to examine the rate of 53BP1-USP28-p53 complex formation in normal or abnormal mitosis as established previously [13], using antibody pairs against 53BP1 and p53 or 53BP1 and USP28 (Fig. 5; data not shown). Cells were treated overnight with RO3306 (RO) to synchronize at the G2-M border and fixed at various timepoints following RO washout and mitotic entry. Examining mitotic cells at time points 10, 20, 30, and 40 minutes after washout revealed that no significant increase in either number or combined area of PLA foci is detectable during normal mitotic divisions (Fig. 4A-B, first panels, and Fig. S4, rows 1-2). However, consistent with prior results, prolonging mitosis by washing out RO in the presence of nocodazole resulted in significant increases in both the number and total area of foci detectable at 80 and 120 minutes (Fig. 4A-B, second panels, and Fig. S4, rows 3-4), though no detectable increases up to 40 minutes after mitotic entry were seen.

**Fig. 4:**
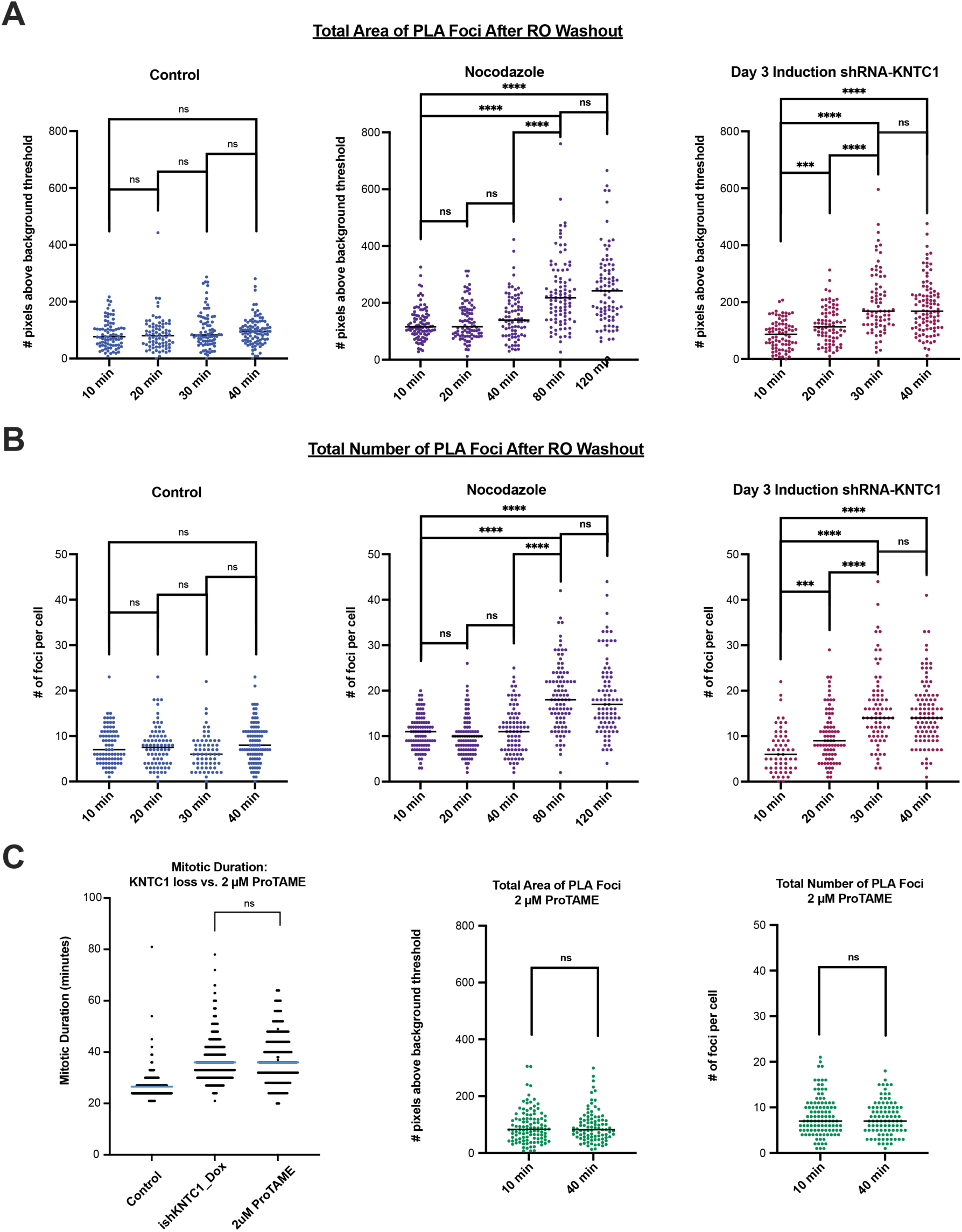
Loss of KNTC1 triggers ectopic formation of 53BP1-USP28-p53 complexes in early mitosis. **(A & B)** Cells arrested at G2 by RO3306 (9.5 μM) were released into mitosis under the indicated conditions for the indicated amounts of time before fixation for PLA analysis of complex formation. In control cells no increase in the total area (**A**) or number (**B**) of PLA foci was seen up to 40 minutes after RO3306 washout (first panel). In the presence of nocodazole (50 ng/mL), an increase in both the area and number of foci was seen between 40 and 80 minutes after washout (second panel) but not before. Following inducible KNTC1 knockdown (Day 3 under doxycycline), an increase in both the area and number of PLA foci was detectable between 10 and 20 minutes after washout, with an additional significant increase between 20 and 30 minutes (third panel). PLA reactions were carried out using primary antibodies against endogenous 53BP1 and p53, *n* > 75, area quantified as the number of pixels above the background threshold, and statistical significances calculated via T-tests with Welch’s correction. Similar PLA results were seen with antibodies against 53BP1 and USP28 (data not shown; see methods). **(C)** Mitotic durations of WT cells treated with low-dose ProTAME (2 μM) resembled those of KNTC1-knockdown cells (left) but did not induce fast formation of 53BP1-USP28-p53 complexes in early mitosis (middle and right). The mean duration in both conditions is ∼36 minutes with no significant difference in distributions (calculated by T-test with Welch’s correction). Cells arrested at G2 with RO3306 (9.5 μM) were released into mitosis in the presence of ProTAME (2 μM) for the indicated amounts of time, fixed, and examined by PLA with primary antibodies against endogenous 53BP1 and p53. No increase in total area (middle) or number (right) of PLA foci was seen up to 40 minutes after RO3306 washout. *n* > 75, statistical significances calculated via T-tests with Welch’s correction.

Strikingly, by examining PLA foci formation after RO washout following knockdown of KNTC1, we saw a significant increase in the number and combined area of foci as early as 10-20 minutes after washout, with an additional increase between 20-30 minutes (Fig. 4A-B, third panels, and Fig. S4, rows 5-6). We further noted that no additional increase is observed between 30 and 40 minutes after washout. Moreover, by releasing WT cells into mitosis in the presence of a low dose of the APC/C inhibitor ProTAME, which slightly slows down mitotic progression, we generated a range of mitotic durations mimicking that of KNTC1 knockout cells (Fig. 4C, first panel). Intriguingly, under this treatment, PLA assays revealed no detectable increase in foci formation up to 40 minutes after mitotic entry (Fig. 4D; Fig. S4, final row), further supporting that accelerated foci formation is specifically caused by loss of KNTC1 rather than slower mitosis. Taken together, these findings demonstrate that in the absence of KNTC1, 53BP1-USP28-p53 complex formation occurs and peaks within 30 minutes after mitotic entry, overlapping with the range of normal mitotic duration, notably faster than other conditions prolonging mitotic durations.

## Discussion

Mitotic integrity is directly linked to the reliability of SAC, a surveillance program ensuring fidelity of chromosome segregation but not its own consistency. The limitation of a system to prove its own consistency from within is a fundamental constraint highlighted by Gödel’s incompleteness theorems [5]. To prove consistency, one must step outside the system. Here, by creating a condition where mitosis operates viably under compromised SAC, we discover that the consistency of fidelity control is safeguarded by a separate system external to mitosis or SAC, the EMS, mediated by 53BP1, USP28, and p53. Mitotic integrity is thus protected by nested layers of surveillance.

We show that mitosis is surveilled by EMS beyond its time efficiency, as KNTC1 loss induces ectopic 53BP1-USP28-p53 complex accumulation over normal mitotic duration. The mechanism by which 53BP1-USP28-p53 complex formation is selectively suppressed or induced during normal, abnormal, early, or late mitosis is unclear [12, 13]. Our finding of an additional input to EMS activation, beyond prolonged mitosis, opens new avenues to dissect the unknown. We conjecture that EMS is activated in response to mitosis operating under suboptimal SAC, a risky condition that not only KNTC1 loss but also prolonged mitosis can face. Indeed, SAC activities are known to fade or weaken during prolonged mitosis even when mitotic errors persist [37]. In extreme cases, mitotic arrest leads to slippage or apoptosis without cell division. In less extreme cases where cells manage to go through delayed mitosis and divide, chromosome segregation may still finish under faded—and thus suboptimal—SAC activities, increasing error risks. The proposed idea is that, instead of taking chances, EMS has evolved in higher animals to invalidate risky mitosis at the system level, regardless of duration. Consistently, in the absence of EMS, KNTC1 loss generates non-population-level cell death at low rates, presumably due to incidental errors accumulated under compromised SAC. Without EMS, cells are “unaware” of SAC consistency, which is likely harmful at animal level; however, such ignorance could perhaps be beneficial to unicellular organisms that survive at all costs, explaining why EMS evolves in animal lineages.

Furthermore, our results reveal that the function of a gene or cellular process can be masked and transcended by external surveillance. EMS imposes SAC consistency as a requirement for cell proliferation, and by doing so, it transcends the function of mitosis from a mechanical machine of cell division to one that also infers cell fitness. In this context, to fully understand how genes or cellular processes work and impact cell behaviors, systematic identifications of external surveillance followed by comparative studies between masked and unmasked conditions at both cellular and animal levels are required.

## Acknowledgments

We are grateful for the *CENP-F^E564P^* cell line kindly provided by the Fava lab. This work was supported by the National Institutes of Health Grant R01 GM-088253 (USA) for MBT.

## Materials and Methods

### Cell Culture

hTERT-RPE1 cells were cultured in DME/F12 (1:1) media supplemented with 10% FBS and 1% penicillin/streptomycin. Puromycin selection was done at 10 mg/mL for 3 days, and blasticidin selection was done at concentration of 5 μg/mL for 5 days. Where indicated, cells were cultured in the presence of drugs: doxycycline (1 μg/mL), BRDU (10 μM), nocodazole (50 ng/mL), centrinone (200 nM), and RO3306 (9.5 μM). HEK-293T cells used for lentiviral production were cultured in DME high glucose media supplemented with 10% FBS and 1% penicillin/streptomycin. Parental transgenic RPE1 lines used in this study include *p53^-/-^, 53BP1^-/-^,* and *USP28^-/-^* cells made previously [9], and *CENP-F^E564P^* cells obtained from the Fava lab [13].

### Individual CRISPR-mediated gene targeting

*CENP-F^E564P^* cells were a generous gift from Luca Fava’s lab [13]. Parental RPE1-Cas9 line and *CENP-F^E564P^*-Cas9 cells were generated in our lab through lentiviral infection carrying the plasmid lentiCas9-Blast (Addgene #52962). Individual CRISPR gRNA sequences were cloned into the lentiGuide-Puro vector (Addgene #52963) or into an altered version of the plasmid swapping the puromycin marker for blasticidin (lentiGuide-Blast) generated in our lab. In this study, we generated the following RPE1 cell lines: *p53^-/-^*, *p53^-/-^; KNTC1^-/+^*, *p53^-/-^; KNTC1^-/-^*, *USP28^-/-^*, *USP28^-/-^; KNTC1^-/+^*, *USP28^-/-^; KNTC1^-/-^*, *53BP1^-/-^*, *53BP1^-/-^; KNTC1^-/+^*, *53BP1^-/-^; KNTC1^-/-^*, and a single heterozygous *KNTC1^-/+^* clone in an otherwise WT genetic background (p53^+/+^*; KNTC1^-/+^*). Sequences of gRNAs used are as follows: p53 (5’-GGTGCCCTATGAGCCGCCTG-3’), KNTC1 (AAACATTCGGAACACTATGG targeting exon25 or ACTCTCCTTCATCAGTCACG targeting exon10), USP28 (TGAGCGTTTAGTTTCTGCAG), and 53BP1 (GTATACCTGCTTGTCCTGTT).

### Genome-wide CRISPR screens

The full genome CRISPR screens were carried out in RPE1 cells of either WT, *p53^-/-^,* or *USP28^-/-^* genetic backgrounds made in the lab [9]. Stable expression of Cas9 was established using the lentiCas9-Blast (Addgene #52962) plasmid. The BRUNELLO pooled gRNA library in the LentiGuide-Puro vector backbone was purchased from Addgene (catalog #73178) and amplified [38]. Viruses were tittered for 40% MOI, and cells were seeded to obtain 100x coverage (8 million infected cells). 24 hours following infection, cells were split into puromycin for 3 days. Following selection, 8 million cells were re-seeded into multilayer flasks for culturing, and the remainder frozen for DNA collection as T_0_. Cells were cultured for 4 more weeks (T_4_), splitting every 3-4 days depending on density. Following extraction of genomic DNA, gRNAs were amplified for sequencing, utilizing the AMPure XP-PCR purification method. Sequencing libraries were generated and sequenced at a depth of 80-100 million reads per sample. The raw counts data are available at GEO, ID # GSE306237 (<u>GEO link</u>). After mapping, read counts per gRNA were normalized to the total read counts at their respective time points. A ratio of reads at T_4_/T_0_ for each gRNA was then used to represent population levels in the exponential growth formula r = [(final population/initial population) ^(1/time)^ −1], resulting in a “percent change in growth” (PCG) value, which is positive or negative for gRNAs enriched or depleted, respectively, at T_4_ vs. T_0_. The gold-standard set of 927 non-essential genes (NEG) was used for normalization [32], with PCG values for each screen shifted to center the mean of non-essential genes at 0. IQR (interquartile range) outlier analysis of all PCGs for NEG was conducted. The upper and lower IQR bounds were used to define neutral regions. Cutoff values for significant enrichment/depletion were then set for each screen, based on the IQR outlier analysis of all PCGs for the NEG. gRNAs for all remaining targeting genes were then analyzed for identification of genes with two or more gRNAs showing selective depletion in the WT screen but a neutral or positive effect within both the *p53^-/-^* and *USP28^-/-^*screens. Gene Ontology (GO) enrichment analysis for cellular components was performed as described [33, 34], using genes with two or more gRNAs selectively depleted in the WT screen as the comparison set.

### KNTC1 mutant genotyping

p53, 53BP1, and USP28 knockout lines were selected based on antibody staining. For clonal isolations with KNTC1 primers, the cut site region was amplified, sequenced, and analyzed with the CRISPREspresso2 analysis tool. KNTC1-exon25 cut site primer sequences: 5’-CCATCTTTCTTTTGGCATCCAGGAAAATACAA-3’ and 5’-GTGCTACACTGCATAGATCCAAG-3’. KNTC1-exon10 cut site primer sequences: 5’-CTCCAGCCTGGGCGACAAGAG-3’ and 5’-CCAGACAAGATACCTGGTACTAACAGAAGGG-3’.

### KNTC1 shRNA

Constructs were designed using the SplashRNA algorithm [35] and expressed from miR-E backbone vectors [36]. Cells were infected with constructs targeting two regions of KNTC1, exons 33 and 55, with the sequences TTCAAATGAAAGAAAGACCTCA and TTTGTATTCTAAACACAGCTCA, respectively.

### Growth Curves analysis

For growth curve analysis, 100,000 cells were seeded in a 6-well dish on Day 0 (with or without the presence of doxycycline) and split alternately every 3 or 4 days thereafter at 1:10 or 1:20 dilutions, respectively, resulting in count values at Day 0, 3, 7, 10, and 14. Total cell counts at time points 7, 10, and 14 were extrapolated by the following formula: [(Cells collected) / (Cells seeded at T_prior_)] • (Total cells at T_prior_).

### Antibodies

The following antibodies were used in this study, with the company, catalog number, species/immunotype, and working concentration used noted in parenthesis: anti-BRDU (BioRad, MCA6144, rat, 1:500), anti-BUBR1 (Abcam, ab6437, mouse IgG 1, 1:400, note – discontinued), anti-Centrin (Millipore-Sigma, 04-1624, mouse IgG 2a, 1:1000), anti-CREST (Immunovision hct-0100, human, 1:50,000), anti-KNTC1 (Santa-Cruz, sc-81853, mouse IgG 2a, 1:50), anti-p21 (Abcam, ab7960, rabbit, 1:200), anti-p53 (Cell Signaling Technology, 9282S, rabbit, 1:400), anti-USP28 (Invitrogen, PA5-52346, rabbit, 1:200), anti-53BP1 (Millipore-Sigma, MAB3802, mouse IgG 1, 1:400), and anti-γH2AX (Millipore-Sigma, 05-636, mouse IgG 1, 1:200). Secondary antibodies with Alexa Fluor™ dyes 488, 594, and 680 were purchased from Invitrogen, all at 1:1000 working concentrations.

### Fixation and Immunofluorescence Staining

Cells grown on glass coverslips were washed with PBS and then fixed with cold methanol at −20°C for 20 minutes before immunostaining. For KNTC1 staining conditions, cells were washed with PBS and followed by 1 minute incubation in PTEM (20 mM PIPES, pH 6.8, 0.2% Triton X-100, 10 mM EGTA, 1 mM MgCl_₂_) for cytoplasmic protein extraction prior to fixation in 4% paraformaldehyde at room temperature for 10 minutes. Fixed cells on coverslips were blocked with 3% BSA and 0.1% Triton X-100 in PBS (PBSBT). Immunofluorescence staining of coverslips with primary and secondary antibodies diluted in PBSBT was carried out at room temperature in light-protected humidity chambers prior to DAPI staining and mounting with ProLong Gold Antifade.

### Proximity-Ligation Assay (PLA)

In situ proximity ligation assay (PLA) is a technique using oligonucleotide-conjugated antibodies to detect single protein-protein interactions within cells by generating specific DNA signals amplified via rolling circle amplification [39]. PLA reactions were done using the reagents and protocol provided by Millipore-Sigma DuoLink. The following two pairs of primary antibodies for 53BP1, USP28, and p53 and their concentrations were used for PLA: (i) rabbit p53 (Cell Signaling, 9282S, 1:400) and mouse 53BP1 (Sigma, MAB3802, 1:400), and (ii) rabbit USP28 (Thermo Fisher, PA5-52346, 1:200) and mouse p53 (Santa Cruz, sc-126, 1:400).

### Microscopy and Image Analysis

Immunostaining images were obtained with an upright microscope (Axis Imager; Carl Zeiss) equipped with Hamamatsu Photonics camera. Images were processed and prepared for figures in AxioVision and ImageJ. Intensity of nuclear p21 was quantified in ImageJ, using the “mean gray value” measurement; the nuclear region was defined based on DAPI and drawn manually. The number of γH2AX foci per nucleus (defined based on DAPI and drawn manually) was quantified in ImageJ. A Gaussian blur with a sigma of 1.0 was applied to each image to minimize false positives from small background variations. For PLA foci, images were first converted to binary using the ImageJ auto local threshold function, following which the number of objects was quantified as the number of foci, and the total area in pixels was measured. For live cell imaging, images were obtained on an inverted microscope (Axiovert; Carl Zeiss), with a 10x objective and a motorized, temperature-controlled stage, in an environmental chamber (Carl Zeiss). Images were obtained with an ORCA Hamamatsu camera and processed in the Zeiss ZEN software. For mitotic duration analyses, mitosis was defined as the first time point at which cells visibly round up through the time point at which either anaphase or cleavage furrow is detectable.

## Supplemental Figure Legends

**Supplemental Fig. S1:**
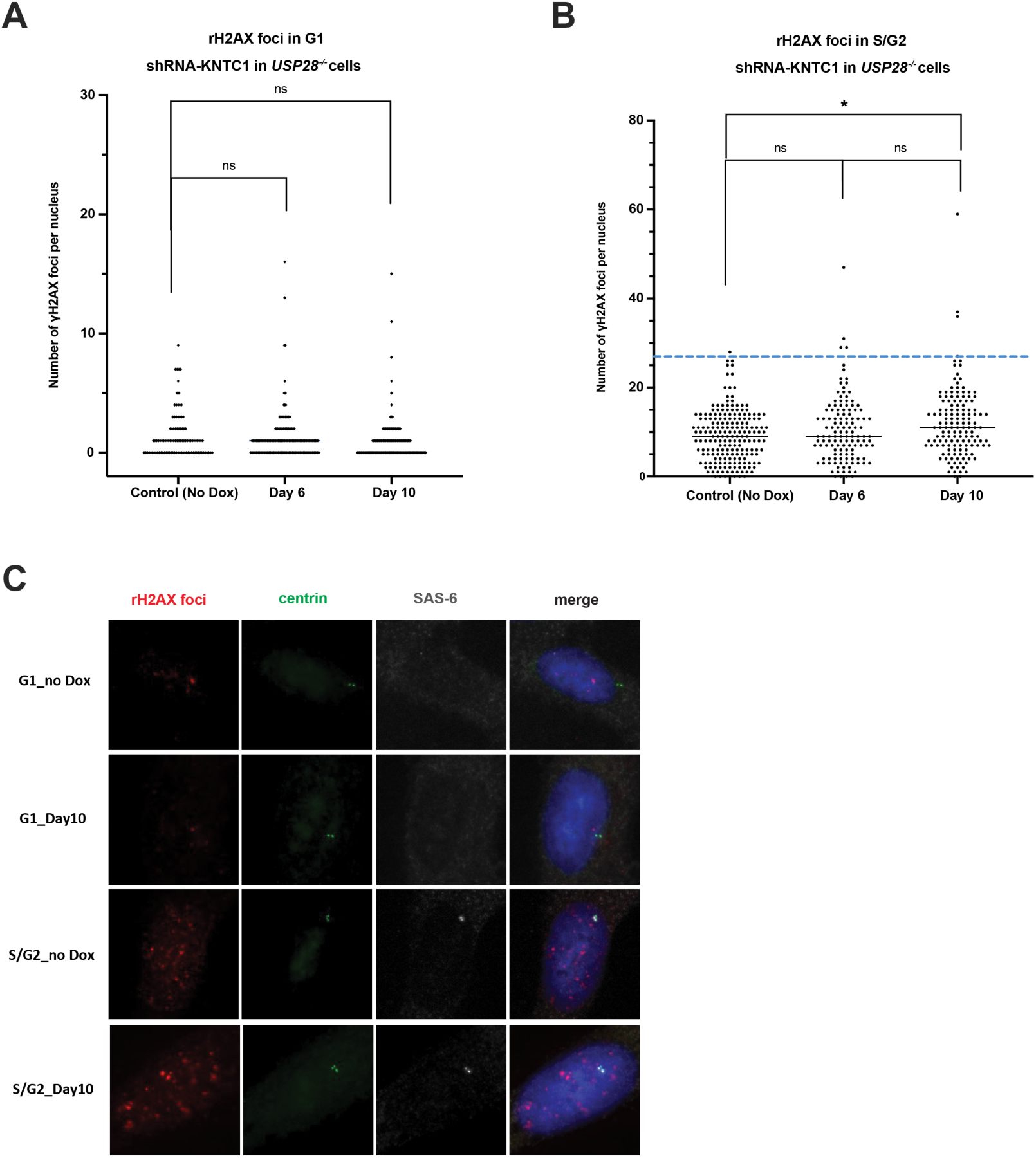
Loss of KNTC1 does not induce DNA damage response. **(A)** Quantification of γH2AX foci per cell in *USP28^-/-^* G1 cells showed no increase following loss of KNTC1. *n* > 60, *N* =2, no statistical difference was found between the two experiments (not shown). Graphs showed combined data, with statistical significance calculated via T-test with Welch’s correction. **(B)** Quantification of γH2AX foci per cell in *USP28^-/-^*cells in S/G2 showed no significant increase following loss of KNTC1 between control and Day 6, or between Day 6 and Day 10. However, a small but significant difference is seen when comparing control to Day 10. Closer examination showed minimal difference in mean number of foci, but the presence of a small number of cells (∼3% at both Day 6 and Day 10) with significantly higher foci counts. *n* > 60, *N* =2, no statistical difference was found between the two experiments (not shown), and graphs showed combined data. Black line at median, blue line at 3 standard deviations above the mean (to control cells), and statistical significance calculated via T-test with Welch’s correction. **(C)** Representative images of γH2AX foci in *USP28^-/-^*control cells (no induction) and at Day 10 following shRNA-KNTC1 induction. Top panels showed G1 cells, identified by the absence of SAS-6 at centrosomes (marked by centrin); bottom panels showed S/G2, identified by the presence of SAS-6 at centrosomes.

**Supplemental Fig. S2:**
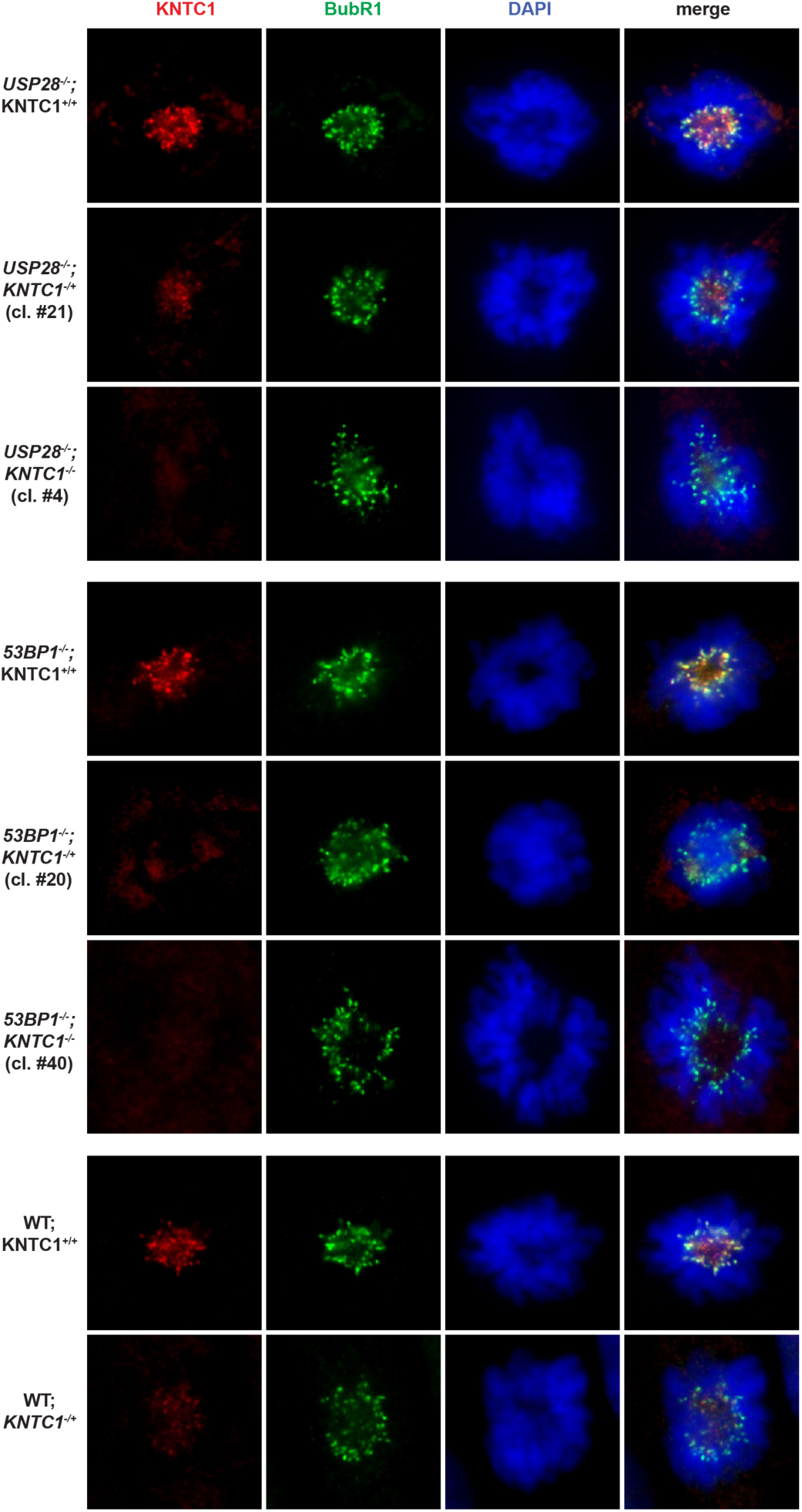
Representative images of KNTC1 staining, following CRISPR targeting and clonal isolation in *USP28^-/-^*, *53BP1^-/-^*, and WT cells. Representative images of normal (KNTC1^+/+^), reduced (*KNTC1^+/-^*), and absent (*KNTC1^-/-^*) KNTC1signal in the *USP28^-/-^*, *53BP1*^-/-^, or WT background as indicated. KNTC1 genotypes were confirmed by sequencing. No *KNTC1^-/-^*clones could be isolated from the WT background.

**Supplemental Fig. S3:**
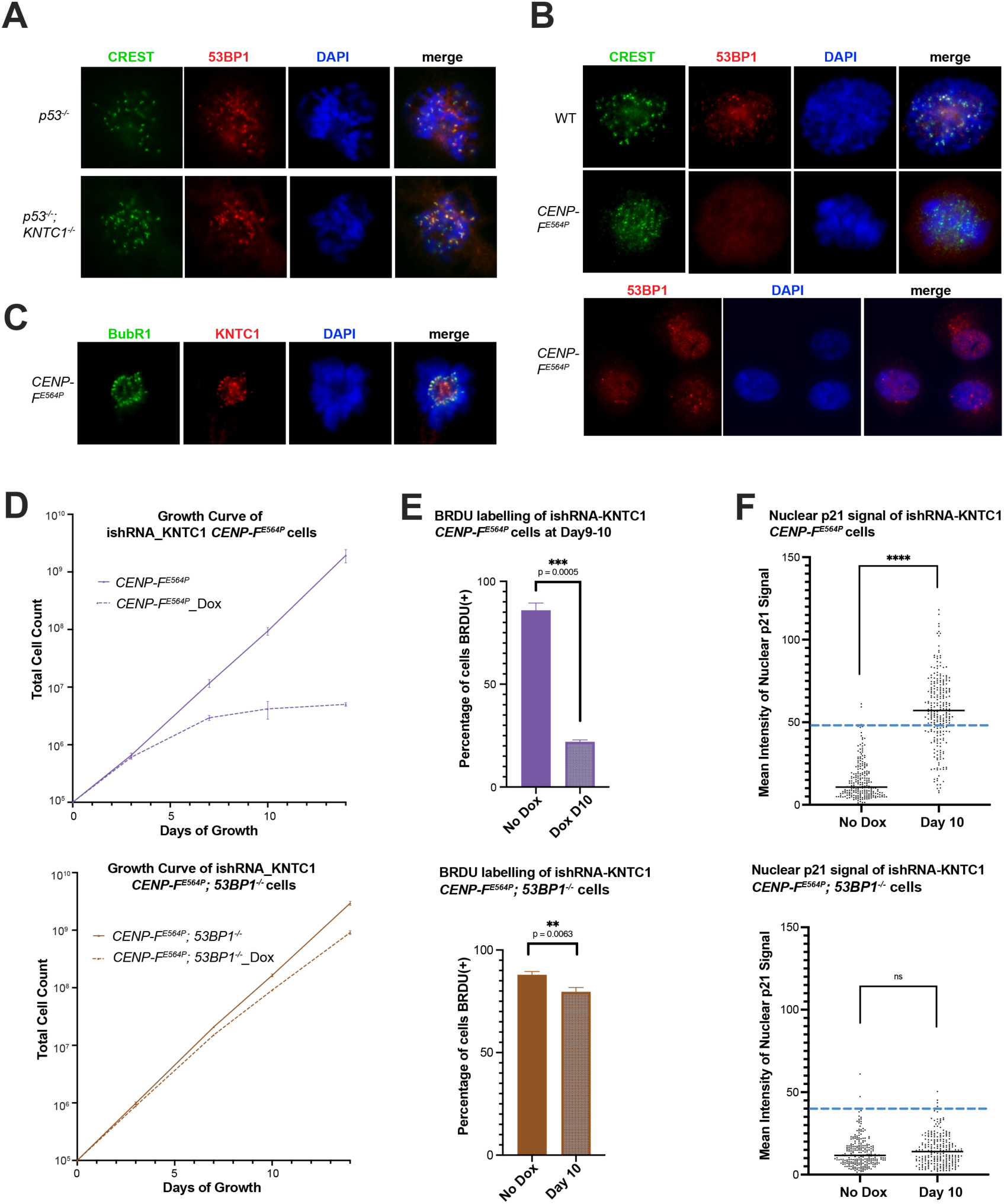
Localizations of 53BP1 and KNTC1 in *KNTC1^-/-^*and *CENP-F^E564P^* cells, and KNTC1 loss-induced cell cycle arrest does not depend on kinetochore 53BP1. **(A)** 53BP1 localizes normally to mitotic kinetochores (CREST) in KNTC1 knockout cells. **(B)** 53BP1 in *CENP-F^E564P^* cells is absent at mitotic kinetochores (top panel) but localized normally to nuclei (bottom panel). **(C)** KNTC1 is localized normally to early mitotic kinetochores (RubR1) in *CENP-F^E564P^* cells. **(D)** Growth curve analyses showed arrest of *CENP-F^E564P^*cells following induction of shRNA-KNTC1 (top). Arrest was rescued in *CENP-F^E564P^;53BP1^-/-^* cells, which continue to grow, despite a slightly slower rate compared to non-induction control (bottom). Data are mean ± SD, *N* = 3, initial cell count = 100,000, and doxycycline (Dox) added at 1 μg/mL. **(E)** 24-hour labeling with 10 μM BrdU from day 9-10 (Day 10) in *CENP-F^E564P^* cells showed nearly 80% of cells stop proliferating by Day 10 (top), a phenotype rescued in the *CENP-F^E564P^;53BP1^-/-^*background (bottom) where cell proliferation continued at the population level, albeit with a slower rate. Doxycycline added at 1 μg/mL for induction, BrdU added 24 hours prior to fixation at a concentration of 10 μM. Data are mean ± SD, *n* > 300, *N* = 3, statistical significance calculated via T-test with Welch’s correction. **(F)** Significant accumulation of nuclear p21 in *CENP-F^E564P^*cells following induction of shRNA-KNTC1, with ∼70% of cells as high outliers to the control cells at Day 10 (top). This phenotype is rescued in *CENP-F^E564P^;53BP1^-/-^*cells, where no significant difference is detected (bottom). Doxycycline, 1 μg/mL. Data are mean ± SD, *n* > 240, black line at median, dashed blue line at 3 standard deviations above mean of control cells, statistical significance calculated via T-test with Welch’s correction.

**Supplemental Fig S4.**
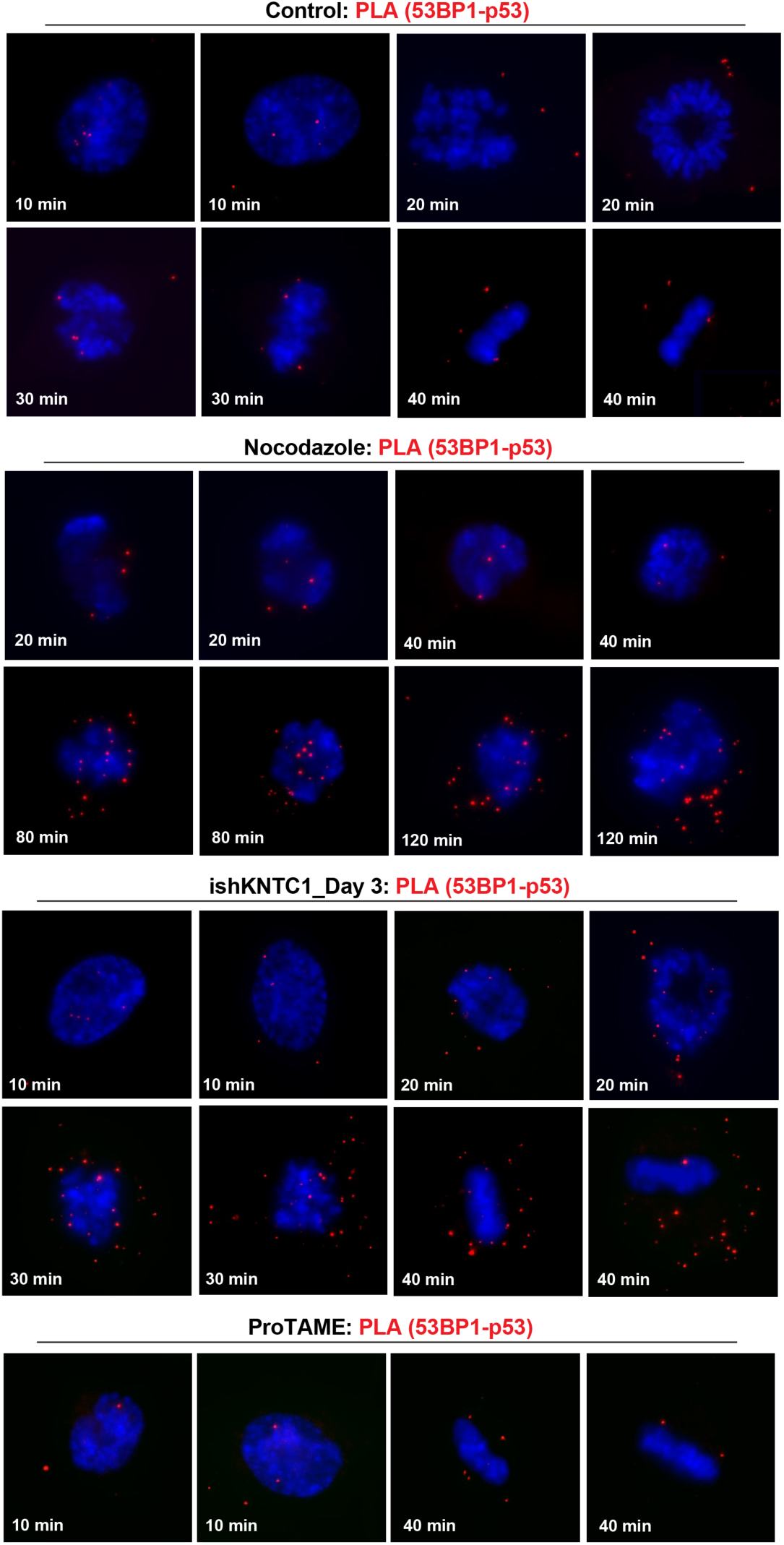
Formation of 53BP1-USP28-p53 complexes revealed by PLA. Representative images of PLA foci using primary antibodies against endogenous p53 and 53BP1 to visualize 53BP1-USP28-p53 complexes in mitosis. Cells were arrested at G2 by RO3306 and released into mitosis under indicated conditions for indicated amounts of time before being fixed and examined by PLA. Cells were released into mitosis without additional drugs (control), in the presence of nocodazole (NOC, 50 ng/mL) or ProTAME (2 μM), or following KNTC1-knockdown (doxycycline 1 μg/mL, Day 3) as indicated.

## Supplemental Table

**Table S1.**

Lists of genes with their corresponding gRNAs negatively impacting WT cell growth while being neutral or positive to *p53^-/-^* or *USP28^-/-^* cells. Neutral impacts were defined and normalized by the "gold-standard" set of 927 non-essential genes.

## Supplemental Movies

**Movie S1**

*p53^-/-^* cells under nocodazole (50 ng/ml). Time-lapse microscopy of asynchronously growing cells at 5-minute intervals.

**Movie S2**

*p53^-/-^; KNTC1^-/+^* cells under nocodazole (50 ng/ml). Time-lapse microscopy of asynchronously growing cells at 5-minute intervals.

**Movie S3**

*p53^-/-^; KNTC1^-/-^* cells under nocodazole (50 ng/ml). Time-lapse microscopy of asynchronously growing cells at 5-minute intervals.

**Movie S4**

Wild-type cells under nocodazole (50 ng/ml). Time-lapse microscopy of asynchronously growing cells at 5-minute intervals.

**Movie S5**

Singleton-Het (p53^WT^; *KNTC1^-/+^*) cells under nocodazole (50 ng/ml). Time-lapse microscopy of asynchronously growing cells at 5-minute intervals.

## Notes

### Competing Interest Statement

The authors have declared no competing interest.

